# Identification of new inhibitors of the toxoplasma gondii by using in-silico drug repurposing

**DOI:** 10.1101/560284

**Authors:** Mosleh Kadkhodamohammadi, Milad Jaberi, Reza Babaei akerdi, Parva Karimimousivandi, Ahmad Ahmadi, Masoud Aliyar

## Abstract

**Introduction:** The common treatment for toxoplasmosis was pyrimethamine. In recent years, it has been found that this parasite is getting resistant to this treatment, therefore urgent alternative treatments are needed.

**Material and Methods:** In this study, by using drug repurposing and in silico methods we tried to make a selective treatment by inhibiting the Calcium-Dependent Protein Kinase 1 from Toxoplasma gondii which doesn’t exist in mammalians. We screened the FDA approved drugs by molecular docking and after ranking them by their binding energies and inspecting the top scored ones, we chose Cefpiramide, Ceftriaxone and Cefotiam as the hit compounds. After that, we used molecular dynamics simulations to test the hit compounds in a much more realistic environment.

**Results:** By analyzing the results, we found that all of the hit compounds and good and can bind strongly to the active site of the protein. Therefore, they can be potential candidates for inhibiting Calcium-Dependent Protein Kinase 1 from Toxoplasma gondii.

**Conclusion:** Moreover, because the predicted compounds are FDA approved drugs, their toxicity profiles are well known and their newly predicted use can be tested in clinical trials.

## Introduction

Toxoplasmosis is an infection disease caused by Toxoplasma gondii and its infection is very widespread. It is estimated that in US and UK, 16 to 40% of the population and in the Central and South America and continental Europe, 50 to 80% of the population are infected (1-6). Toxoplasma gondii is a dangerous opportunistic pathogen and invade the entire body, thus effective treatments are urgent.

To this date, there are only a few treatments for Toxoplasma gondii infection: sulfadiazine and pyrimethamine or clindamycin, that have severe side effects. Therefore, effective drugs with good efficacy and little side-effects are needed.

In 2010, a study (7) proved that calcium-dependent protein kinase 1 in Toxoplasma gondii is a selective target to inhibit the growth and indeed the invading of this parasite. By designing effective compounds with great efficacy, they managed to overcome Toxoplasma gondii. Moreover, since this target is not present in human beings, it is a perfect target for this treatment.

According to the Tufts Center for the Study of Drug Development (CSDD), the high cost of newly approved drugs is due to the costly R&D (Research and Development) processes. It shows that for bringing a new approved drug to the market, it takes 15 years and 2 billion dollars on average (8-10) and moreover, risky drugs such as antibiotics that can get resistant to treatment in a matter of short time, attract less attention from pharmaceutical companies (11, 12). A good solution to this problem is drug repurposing.

Drug repurposing is using an old drug for a new indication (13-18). Drug repurposing is a famous and arguable method to bring down the cost of research and development in drug discovery. Health organizations such as the Food and Drug Administration of USA (FDA) examine and evaluate new chemical entities for a certain indication in preclinical and clinical trials. They make sure that the drug is safe and has the efficacy to treat an specific disease and then approve it to be used in public (19). However, if an old drug is a candidate for another indication, it does not need to pass any safety tests and all of the pharmacokinetics and safety profiles are available (20).

Wet lab experiments, sometimes, can cost large amounts of money and using computational methods, as well as drug repurposing, bring down expenses in experiments (21, 22). Computational methods in drug discovery such as Ligand-based or structure-based drug designing are gaining attention and numerous projects have been carried out over the last years (23-27). In the past years there have been a few studies that tried to find a treatment for Toxoplasma gondii by in silico methods and they have chosen profilin and dihydrofolate reductase as their target protein (28, 29). However, these targets might not be safe targets to inhibit and more selective treatments are needed.

In this study, by using drug repurposing and computational drug design methods such as Virtual High Throughput Screening, Molecular docking, Molecular dynamics simulation and related analysis, we ought to discover new potential compounds that can inhibit the Calcium-Dependent Protein Kinase 1 from Toxoplasma gondii in order to treat this parasite.

## Materials and Method

### Obtaining the structures

The crystallographic structure of Calcium-Dependent Protein Kinase 1 from Toxoplasma gondii (TgCDPK1) (PDB code: 3I7B in complex with NM-PP1) (7) was obtained from Protein Data Bank (30). We deleted the co-crystalized ligand (NM-PP1) and the crystallographic water molecules. The missing segments in the structure were built by MODELLER software (31)

The structures of the FDA approved drugs were obtained from Drug Bank database (32). The hydrogens were added and the charges were assigned by using Gasteiger–Marsili method (33). These modifications were done by the UCSF chimera software (34). We also used UCSF chimera for visualization and validation of the structures. We only included compounds with a molecular weight of between 100 and 800 Da in our data-set.

### Virtual screening

For the virtual screening (35), we used molecular docking algorithm using Ledock software package (36). We prepared the protein via Lepro tool from Ledock package. Then, we defined a cubic box in the center of the active site of TgCDPK1with 20*20*20 Å diameters. We performed the virtual screening via Ledock docking program and then ranked the docked poses according to their binding Energies. Then we chose our hit compounds from the top scored compounds. We used NM-PP1 as the reference ligand since it is a proved inhibitor of Calcium-Dependent Protein Kinase 1.

### Molecular Dynamics simulations

The topology files and the parameters of the potential compounds were generated by ACPYPE tool (37). For the MD simulations, we used GROMACS 2018 software (38) and Amber99SB force field (39) and TIP3P water mode. We chose triclinic boxes for the simulation systems and the boxes were under periodic boundary conditions (PBCs). the complexes consisted of the Calcium-Dependent Protein Kinase 1 enzyme and the docked poses of the hit compounds in the active site of the enzyme. The complexes were placed in the middle of the simulation boxes. The gap between the complex and the box edges was set to 1 Å. The boxes were filled with water and 150 mM of NaCl ions were added to neutralize the systems. The simulation systems were energy minimized using the steepest descent minimization algorithm. Then 100 ps NVT (constant number of particles (N), volume (V), and temperature (T)) and 300 ps NPT (constant number of particles (N), pressure (P), and temperature (T)) equilibrations were performed for each system. For the MD production, we simulated all the systems for 10 ns.

The MD simulation were performed to make sure that the identified hit compounds are not some noises in our data and in order to evaluate them we used analyzes such as Root Mean Square Distance (RMSD), Root Mean Square Fluctuation (RMSF), the number of hydrogen bonds and interaction diagrams to validate our findings.

## Results & Discussion

### Virtual screening

We screened roughly 1800 FDA approved drug in order to fine appropriate compounds that can bind to and also inhibit the Calcium-Dependent Protein Kinase 1. We supposed that this strategy can be effective since this enzyme is one of the key elements in Toxoplasma gondii’s invasion. By using the FDA approved drugs as the initial data set for the virtual screening, we wanted to find safe compounds, which have been widely used and extensively tested, that can be used as a treatment for Toxoplasmosis.

After the virtual screening, we ranked the docked poses by their interaction energies. We analyzed the top 20 compounds visually and inspected them in the binding site of the target enzyme and eventually, we chose 3 compounds for further calculations (indicated in red). As it is shown in the table 1, amazingly, in the top 20 scored compounds, 7 compounds were actual kinase inhibitors (indicated in purple). This means that the docking procedure has been done very well and accurately and also it shows that there is a good possibility for the predicted compounds to be effective inhibitors. Additionally, in order to validate the docking procedure, we also docked the co-crystallized ligand. The interaction energy of the co-crystallized ligand was −7.03 kcal/mol which is quite good and reasonable.

**Table 1.**
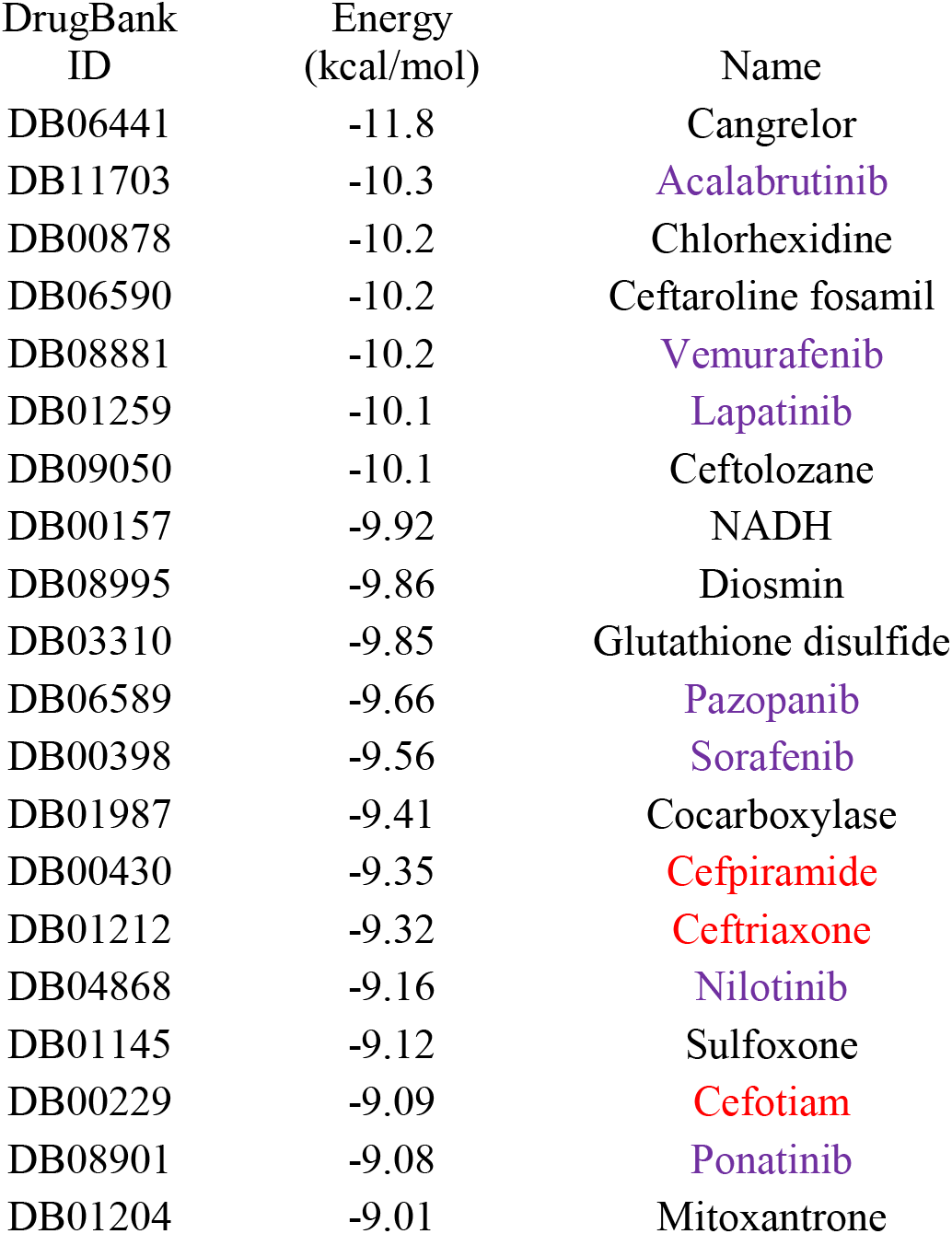
The ranking of the top 20 scored compounds in order of their binding energy (Kcal/mol). The hit compounds are indicated in red and the kinase inhibitors are indicated in purple.

### Molecular dynamics simulation

We took Cefpiramide, Ceftriaxone and Cefotiam as the hit compounds and to test them in molecular dynamics simulation. In a molecular dynamics simulation, environmental conditions such as temperature, pressure, solvent and ion molecules are present. These conditions mimic the real world in the cell as much as possible. The movements of the compounds in the active site of the enzyme can help us understand the key residues and key interactions which are essential in the binding mechanism.

To further investigate the binding mechanism of the hit compounds, we performed 4 simulations. We simulated each of the hit compounds plus the co-crystallized ligand for 10 ns. After the production runs of 10 ns, we validate the runs by RMSD, RMSF calculations.

RMSD stands for Root Mean Square Distance and it is used to measure the movement of the simulated structure according to the initial structure. In a molecular dynamics simulation, it is very important for the structure to reach an equilibration state and get stable. In a RMSD plot, at the very first moments, the value of RMSD of the simulated structure suddenly rise and it is because of the sudden change in the structure. This change in the structure is mostly related to the crystallizing method. In the crystallization, the protein loses its motion and become almost rigid and when it is put in a box full of solvent and ion molecules, it gains its motion back and therefore the structure has little changes. As it is shown in the Fig. 1, the value of RMSD of all systems reach an equilibration state except for Ceftriaxone. In about 6 ns, the protein structure starts to change and as a result of that, the value of RMSD goes up. However, by analyzing the trajectory movie, we found that this rise was due to the conformational changes of Ceftriaxone in the binding site of the protein and it shows that more production run is needed to get a correct conformation and orientation for this ligand. Despite this, the conformation of this compound was acceptable and reasonable and had good interactions with the residues in the protein which will be discussed later.

**Figure 1.**
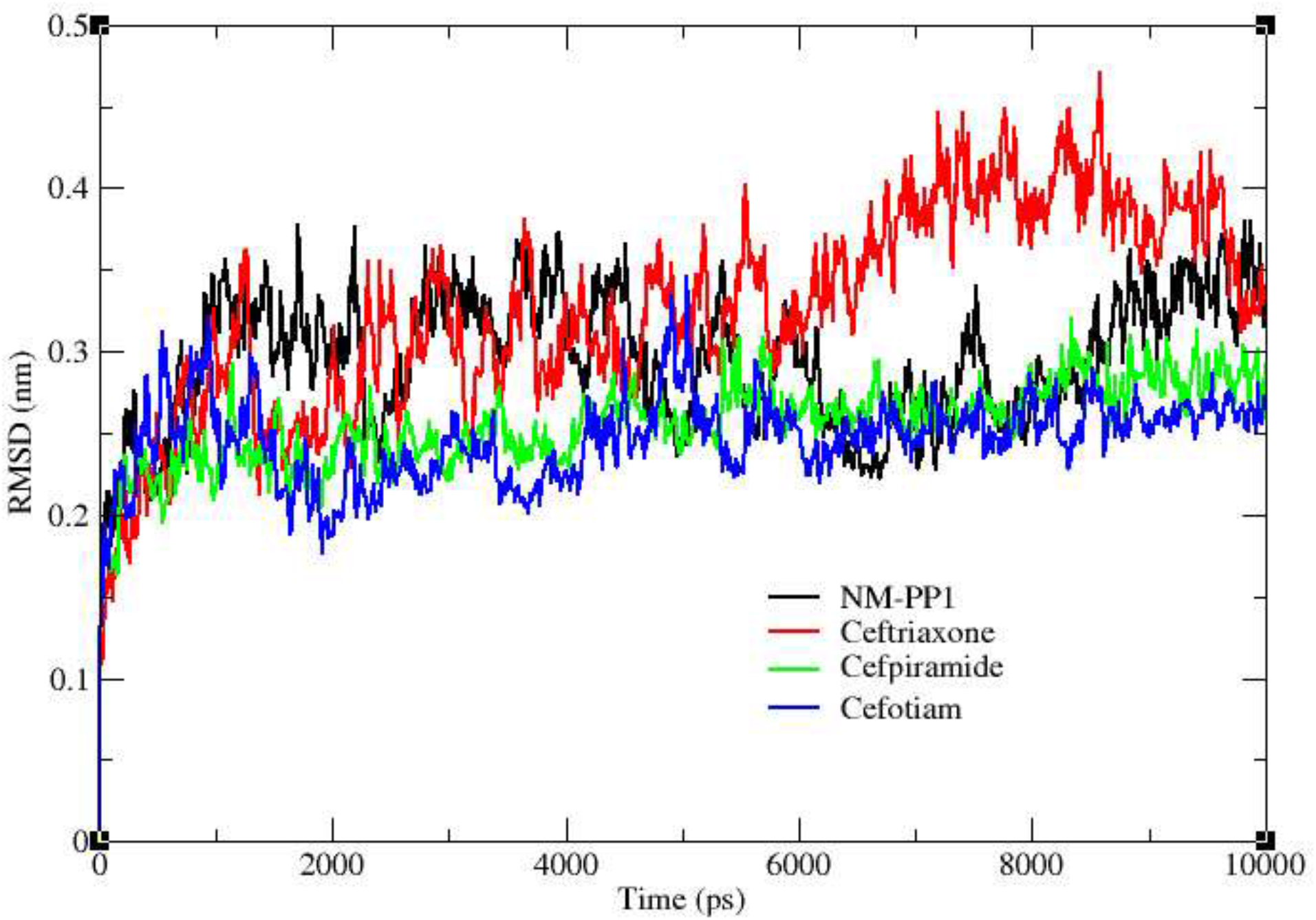
Backbone RMSD values of protein-ligand complexes in the 10-ns period of MD simulations.

RMSF stands for Root Mean Square Fluctuation and it shows the average movement of each residue during the simulation period. In the Calcium-Dependent Protein Kinase 1 from Toxoplasma gondii, the gatekeeper position of the active site is owned by the Gly128. This position is critical in the ligand binding and in fact, this is the main difference between this kinase and mammalian kinases (7) and this allows us to design or find selective inhibitors that have the least side effects in the human body.

The N-terminal and the C-terminal of the proteins always have high amount of fluctuations and most of the time their movement are not considerably important. As it is shown in the Fig. 2, the RMSF plot, all the compounds made the residues have the same fluctuation as the co-crystallized ligand except for the ceftriaxone. As it was mentioned in the RMSD plot, the Ceftriaxone complex need more production run. Because, in theory, we think it is jiggling in the active site to find the best orientation and because of that, it makes some of the region fluctuate more that usual. However, other compounds are considered good and their RMSF values shows that they can manage the residues fluctuations just like the co-crystallized ligand which is a good inhibitor.

**Figure 2.**
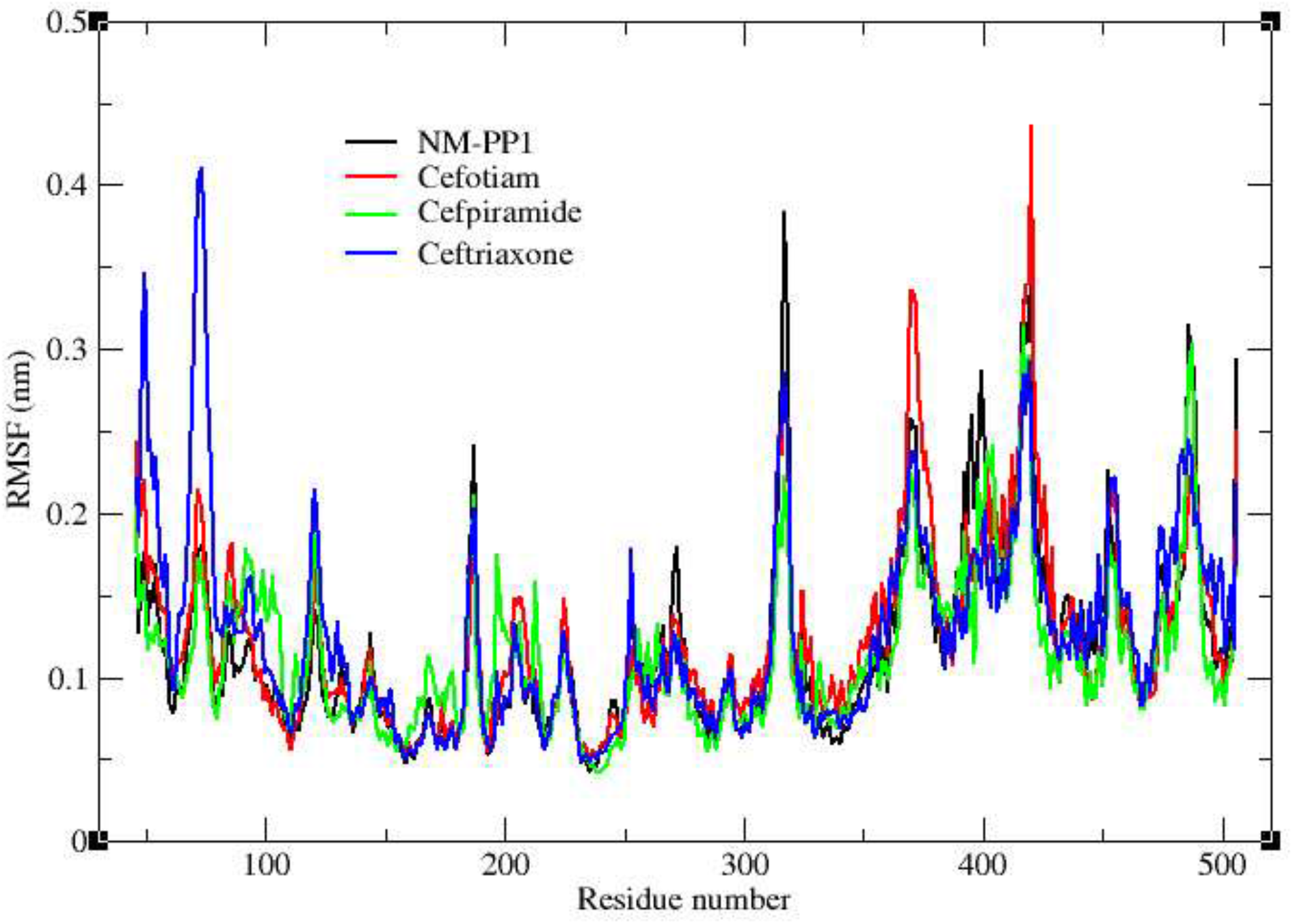
RMSF values of backbone atoms against residue number of protein-ligand complexes.

Monitoring the interactions between the compounds and the residues inside the binding site is extremely important. These interactions widen our point of view about the properties of the binding pocket. The most important interactions are the strongest ones. One of the strongest interactions in a protein-ligand complex is the hydrogen bond. Monitoring the position and indeed the number of hydrogen bonds during the simulation period can help us find the correct conformation and orientation of the ligand inside the binding site. The number of the formed hydrogen bonds between each ligand and the residues inside the binding pocket during the simulation is shown in Fig. 3. As it is shown in the Fig. 3, the number of the hydrogen bonds of the hit compounds is much more than the co-crystallized ligand and this indicate that the hit compounds have a good possibility to be good binders.

**Figure 3.**
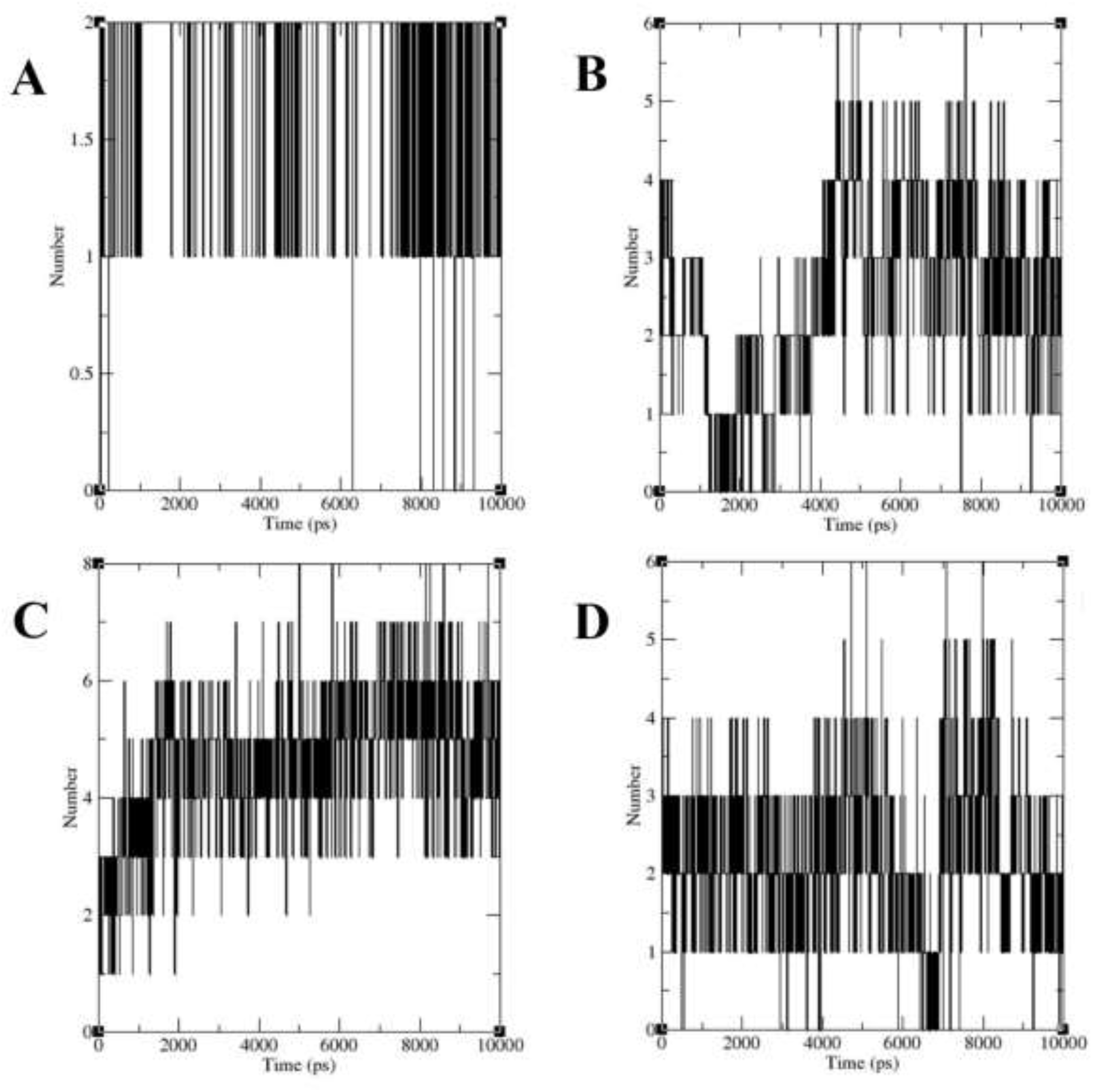
The number of hydrogen bonds between the hit compounds and the residues of the active site of Calcium-Dependent Protein Kinase 1 from Toxoplasma gondii (TgCDPK1). A) TgCDPK1-NMPP1, B) TgCDPK1-Cefotiam, C) TgCDPK1-Cefpiramide, D) TgCDPK1-Ceftriaxone.

Visualization of the ligand in the binding can help us understand the properties of the binding site and also help us find the critical interactions in the ligand’s binding mechanism. Therefore, we inspected the ligand in the binding site via the 2D interaction diagrams. The 2D interaction diagrams are shown in the Fig. 4. As it is shown in the Fig. 4, the number and also the strength of the interactions of the hit compounds are more than the co-crystallized ligand which indicate that the hit compounds are good binders and can strongly attach to the binding site.

**Figure 4.**
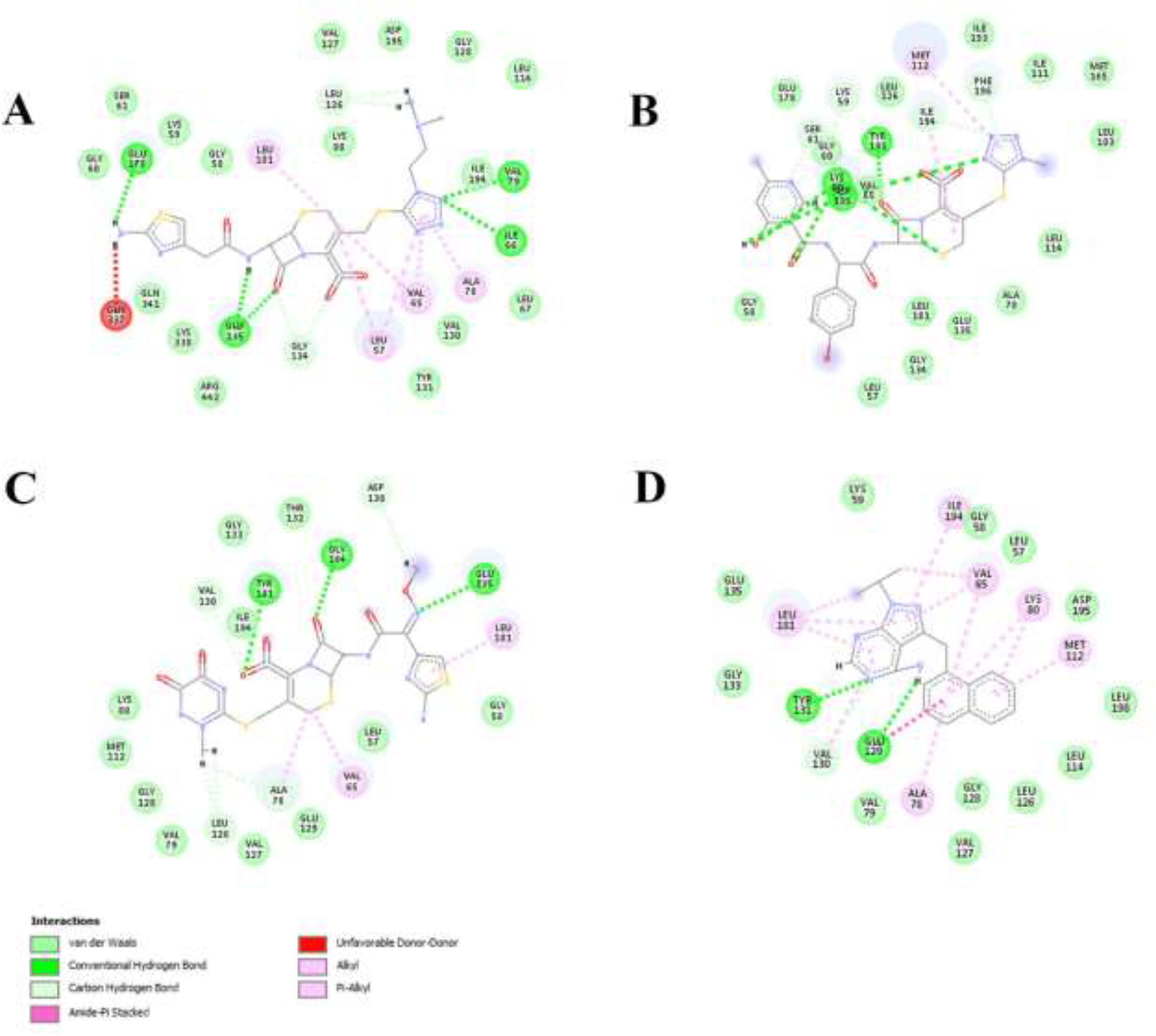
The 2D interaction diagrams of the hit compounds in the active site of the TgCDPK1. A) TgCDPK1-Cefotiam, B) TgCDPK1-Cefpiramide, C) TgCDPK1-Cefriaxone, D) TgCDPK1-NMPP1.

Moreover, by inspecting the 2D diagrams, we can find the key residues in the binding of the hit compounds. We found that in cefotiam-protein complex, GLU178, GLU135, ILE66 and VAL79 formed hydrogen bonds and LEU181, LEU57, VAL65 and ALA78 formed pi-Alkyl interactions and other residues had van der Waals interactions with the ligand. In Cefpiramide-protein complex, TYR131, LYS80 and ASP195 formed hydrogen bonds and PHE196 formed a Pi-Pi interaction and MET112 and ILE194 formed pi-Alkyl interactions. In ceftriaxone-protein complex, TYR131, GLU135 and GLY134 formed hydrogen bonds and LEU181 formed a pi-Alkyl interaction. In the NM-PP1-protein complex, TYR131 and GLU129 formed hydrogen bonds. Since NM-PP1 has four aromatic rings, several residues formed pi-Alkyl interactions and also GLU129 formed an amide-Pi stacked interaction.

The overall analysis showed that the hit compounds are quite good and can strongly interact and bind to the residues in the active site of the Calcium-Dependent Protein Kinase 1.

## Conclusion

In the last decade, drug discovery has become a major concern due to the lack of efficient treatments and the increasing incidence of bacterial and tropical diseases in the world. The reason for the lack of efficient treatments is the complexity of drug design and also the huge cost of clinical trials. In order to get around these issues, we decided to choose drug repurposing as our course of study. In drug repurposing, by finding new uses for old and sheltered drugs, you can simply pass most of the safety test in the clinical trials. This can save a lot of time and money and can make drug discovery extremely efficient.

Toxoplasmosis in a wide spread disease in the world and urgent treatment are needed. In this study by using drug repurposing and in silico methods such as molecular docking and molecular dynamics simulations we tried to find good inhibitors for Calcium-Dependent Protein Kinase 1 from toxoplasma gondii. This enzyme is one the key enzymes in the invasion mechanism of this creature and does not exist in mammalians. Therefore, it is a perfect target for a selective treatment. We used FDA approved drugs as our data set for the virtual screening and after ranking them by their binding energies and inspecting the top scored compounds, we chose the hit compounds. After that, we simulated the hit compounds to get a dynamic view of their binding and by analyzing the results we found that the hit compounds are quite good at binding to the active site and therefore can be potential inhibitors for this enzyme.

Since the main idea of our study is drug repurposing, if any of these predicted drugs may actually inhibit this enzyme, there is no barriers for testing them in phase 2 clinical trials.

## Acknowledgment

We want to thank Professor Aryapour for kind advices about the methods. We also want to thank Mr. Farzin Sohraby, the vice president of Sabzeh Bahonar NGO, for funding this project.

## References

1. Hill D, Dubey JP. Toxoplasma gondii: transmission, diagnosis and prevention. Clinical Microbiology and Infection. 2002;8(10):634–40.

2. Furtado JM, Smith JR, Belfort R, Jr., Gattey D, Winthrop KL. Toxoplasmosis: a global threat. J Glob Infect Dis. 2011;3(3):281–4.

3. Wang ZD, Wang SC, Liu HH, Ma HY, Li ZY, Wei F, et al. Prevalence and burden of Toxoplasma gondii infection in HIV-infected people: a systematic review and meta-analysis. Lancet HIV. 2017;4(4):e177–e88.

4. Flegr J, Prandota J, Sovickova M, Israili ZH. Toxoplasmosis--a global threat. Correlation of latent toxoplasmosis with specific disease burden in a set of 88 countries. PLoS One. 2014;9(3):e90203.

5. Wilking H, Thamm M, Stark K, Aebischer T, Seeber F. Prevalence, incidence estimations, and risk factors of Toxoplasma gondii infection in Germany: a representative, cross-sectional, serological study. Scientific Reports. 2016;6:22551.

6. Jones JL, Kruszon-Moran D, Wilson M, McQuillan G, Navin T, McAuley JB. Toxoplasma gondii Infection in the United States: Seroprevalence and Risk Factors. American Journal of Epidemiology. 2001;154(4):357–65.

7. Ojo KK, Larson ET, Keyloun KR, Castaneda LJ, Derocher AE, Inampudi KK, et al. Toxoplasma gondii calcium-dependent protein kinase 1 is a target for selective kinase inhibitors. Nat Struct Mol Biol. 2010;17(5):602–7.

8. DiMasi JA, Hansen RW, Grabowski HG. The price of innovation: new estimates of drug development costs. J Health Econ. 2003;22(2):151–85.

9. Morgan S, Grootendorst P, Lexchin J, Cunningham C, Greyson D. The cost of drug development: a systematic review. Health Policy. 2011;100(1):4–17.

10. Dickson M, Gagnon JP. Key factors in the rising cost of new drug discovery and development. Nat Rev Drug Discov. 2004;3(5):417–29.

11. Conly J, Johnston B. Where are all the new antibiotics? The new antibiotic paradox. Can J Infect Dis Med Microbiol. 2005;16(3):159–60.

12. Simpkin VL, Renwick MJ, Kelly R, Mossialos E. Incentivising innovation in antibiotic drug discovery and development: progress, challenges and next steps. J Antibiot (Tokyo). 2017.

13. Novac N. Challenges and opportunities of drug repositioning. Trends Pharmacol Sci. 2013;34(5):267–72.

14. Iorio F, Bosotti R, Scacheri E, Belcastro V, Mithbaokar P, Ferriero R, et al. Discovery of drug mode of action and drug repositioning from transcriptional responses. Proc Natl Acad Sci U S A. 2010;107(33):14621–6.

15. Tan F, Yang R, Xu X, Chen X, Wang Y, Ma H, et al. Drug repositioning by applying ‘expression profiles’ generated by integrating chemical structure similarity and gene semantic similarity. Mol Biosyst. 2014;10(5):1126–38.

16. Ashburn TT, Thor KB. Drug repositioning: identifying and developing new uses for existing drugs. Nat Rev Drug Discov. 2004;3(8):673–83.

17. Harrison C. Signatures for drug repositioning. Nat Rev Genet. 2011;12(10):668.

18. Jin G, Wong ST. Toward better drug repositioning: prioritizing and integrating existing methods into efficient pipelines. Drug Discov Today. 2014;19(5):637–44.

19. What does “FDA approval” mean? [Available from: http://www.rotlaw.com/legal-library/what-does-fda-approval-mean/.

20. Gupta SC, Sung B, Prasad S, Webb LJ, Aggarwal BB. Cancer drug discovery by repurposing: teaching new tricks to old dogs. Trends Pharmacol Sci. 2013;34(9):508–17.

21. Quinn TA, Kohl P. Combining wet and dry research: experience with model development for cardiac mechano-electric structure-function studies. Cardiovasc Res. 2013;97(4):601–11.

22. Penders B, Horstman K, Vos R. Walking the Line between Lab and Computation: The “Moist” Zone. BioScience. 2008;58(8):747–55.

23. Sliwoski G, Kothiwale S, Meiler J, Lowe EW, Jr. Computational methods in drug discovery. Pharmacol Rev. 2014;66(1):334–95.

24. Leelananda SP, Lindert S. Computational methods in drug discovery. Beilstein Journal of Organic Chemistry. 2016;12:2694–718.

25. Katsila T, Spyroulias GA, Patrinos GP, Matsoukas M-T. Computational approaches in target identification and drug discovery. Computational and Structural Biotechnology Journal. 2016;14(Supplement C):177–84.

26. Hodos RA, Kidd BA, Shameer K, Readhead BP, Dudley JT. In silico methods for drug repurposing and pharmacology. Wiley Interdiscip Rev Syst Biol Med. 2016;8(3):186–210.

27. Vanhaelen Q, Mamoshina P, Aliper AM, Artemov A, Lezhnina K, Ozerov I, et al. Design of efficient computational workflows for in silico drug repurposing. Drug Discovery Today. 2017;22(2):210–22.

28. Pacheco Homem D, Flores R, Jr., Tosqui P, de Castro Rozada T, Abicht Basso E, Gasparotto A, Jr., et al. Homology modeling of dihydrofolate reductase from T. gondii bonded to antagonists: molecular docking and molecular dynamics simulations. Mol Biosyst. 2013;9(6):1308–15.

29. Kadirvel P, Anishetty S. Potential role of salt-bridges in the hinge-like movement of apicomplexa specific beta-hairpin of Plasmodium and Toxoplasma profilins: A molecular dynamics simulation study. J Cell Biochem. 2018;119(4):3683–96.

30. Berman HM, Westbrook J, Feng Z, Gilliland G, Bhat TN, Weissig H, et al. The Protein Data Bank. Nucleic Acids Res. 2000;28(1):235–42.

31. Eswar N, Webb B, Marti-Renom MA, Madhusudhan MS, Eramian D, Shen M-y, et al. Comparative Protein Structure Modeling Using Modeller. Current protocols in bioinformatics / editoral board, Andreas D Baxevanis [et al]. 2006;0 5:Unit-5.6.

32. Wishart DS, Knox C, Guo AC, Shrivastava S, Hassanali M, Stothard P, et al. DrugBank: a comprehensive resource for in silico drug discovery and exploration. Nucleic Acids Res. 2006;34(Database issue):D668–72.

33. Gasteiger J, Marsili M. Iterative partial equalization of orbital electronegativity—a rapid access to atomic charges. Tetrahedron. 1980;36(22):3219–28.

34. Pettersen EF, Goddard TD, Huang CC, Couch GS, Greenblatt DM, Meng EC, et al. UCSF Chimera--a visualization system for exploratory research and analysis. J Comput Chem. 2004;25(13):1605–12.

35. Muegge I, Oloff S, Abraham DJ. Virtual Screening. Burger’s Medicinal Chemistry and Drug Discovery: John Wiley & Sons, Inc.; 2003.

36. Ledock. Ledock [wwwlepharcom](http://www.lephar.com).

37. Sousa da Silva AW, Vranken WF. ACPYPE - AnteChamber PYthon Parser interfacE. BMC Res Notes. 2012;5:367.

38. Pronk S, Páll S, Schulz R, Larsson P, Bjelkmar P, Apostolov R, et al. GROMACS 4.5: a high-throughput and highly parallel open source molecular simulation toolkit. Bioinformatics. 2013;29(7):845–54.

39. Hornak V, Abel R, Okur A, Strockbine B, Roitberg A, Simmerling C. Comparison of multiple Amber force fields and development of improved protein backbone parameters. Proteins: Structure, Function, and Bioinformatics. 2006;65(3):712–25.

